# Ventilation-induced epithelial injury drives biological onset of lung trauma *in vitro* and is mitigated with anti-inflammatory therapeutics

**DOI:** 10.1101/2021.05.02.441992

**Authors:** Eliram Nof, Arbel Artzy-Schnirman, Saurabh Bhardwaj, Hadas Sabatan, Dan Waisman, Ori Hochwald, Maayan Gruber, Liron Borenstein-Levin, Josué Sznitman

## Abstract

Mortality rates among patients suffering from acute respiratory failure remain perplexingly high despite maintenance of blood homeostasis. The *biotrauma* hypothesis advances that mechanical forces from invasive ventilation trigger immunological factors that spread systemically. Yet, how these forces elicit an immune response remains unclear. Here we show that flow-induced stresses under mechanical ventilation can injure the bronchial epithelium of ventilated *in vitro* upper airway models and directly modulate inflammatory cytokine secretion associated with pulmonary injury. We identify site-specific susceptibility to epithelial erosion in airways from jet-flow impaction and measure an increase in cell apoptosis and modulated secretions of cytokines IL-6, 8 and 10. We find that prophylactic pharmacological treatment with anti-inflammatory therapeutics reduces apoptosis and pro-inflammatory signaling during ventilation. Our 3D *in vitro* airway platform points to a previously overlooked origin of lung injury and showcases translational opportunities in preclinical pulmonary research towards protective therapies and improved protocols for patient care.

## Introduction

Acute respiratory failure epitomizes a deleterious lung condition associated with high mortality rates (>35%) in critical care patients undergoing ventilatory support^1^, despite the maintenance of blood homeostasis^2,3^. In deciphering this conundrum, the pulmonary *biotrauma* theory suggests a multi-factorial cascade, starting with invasive mechanical forces triggering the release of immunological mediators in the lungs^4^. Cyclic respiratory airflows are hypothesized to facilitate the distribution of such mediators and their translocation across the alveolar-capillary barrier into the wider systemic circulation^2^. In turn, inflammatory effects may spread and amplify throughout the body leading to multi-organ failure and eventually death^2^.

Biotrauma was first proposed in the context of invasive mechanical ventilation; a life supporting clinical intervention also recognized to concurrently cause or worsen lung morbidity^3^. Most recently, mechanical ventilation has gained increased scrutiny amidst the COVID-19 pandemic^5–7^, owing to alarmingly higher mortality rates among patients requiring respiratory support^8,9^ (∼50% to 97%). The main physical mechanisms identified as contributing to pulmonary injury^10^ include the overstretching during ventilation of lung tissue from over inflation (known as *volu*- or *baro*trauma) and the repeated opening and collapse of small airway units at excessively low volumes (i.e., *atelec*trauma). Various protective ventilation protocols have emerged to marginally improve patient outcomes and reduce mortality^11^. Yet eliminating injurious mechanical forces is elusive as underlined in clinical trials^12^. Recent efforts have begun exploring therapeutic opportunities (e.g. gene delivery) to mitigate lung injury in the deep alveolar regions^13^. Still, it remains widely unknown to what extent mechanical forces influence the downstream precursors leading to biotrauma. Indirect evidence from animal models^14,15^ and clinical studies^11,16^ has provided seminal support for the biotrauma hypothesis. However, a direct corroboration *in vivo* is challenging since lung-derived inflammatory biomarkers in patients are not readily collected in clinical settings in detecting the origins of cytokine release and trafficking^17^.

To bridge this gap, engineered *in vitro* lung models leveraging advances in tissue engineering and microfabrication have been increasingly utilized for advancing preclinical pulmonary research and explore lung injury^18,19^. For example, *in vitro* studies have characterized wounding in alveolar epithelial cells subjected to cyclic over-stretching, including the release of inflammatory cytokines, impaired structural integrity via changes in tight junctions and plasma membrane breaks, and increased incidence of apoptosis^20–22^. Cellularized *in vitro* models have typically focused on the airway epithelial barrier of the deep acinar regions^23^, where mechanical stresses are dominated by tensile strains when the alveolar airway barrier expands and contracts cyclically^24^. In contrast, there has been little emphasis on the lungs’ proximal regions (i.e., upper airways), where airflow dynamics are most prominent. Namely, the exposure of bronchial epithelial cells to respiratory flow-induced shear stresses has been proposed as a potential link between large ventilation pressures and morbidity and mortality^25^; a situation strongly correlated in ventilated patients undergoing surgery in the absence of prior lung injury^26^. In support, computational fluid mechanics (CFD) based *in silico* studies have found that upper airway flow phenomena and ensuing wall shear stresses (WSS) may contribute to ventilator-induced lung injury (VILI) via the biotraumatic pathway^27–29^. We have also recently underlined the plausible occurrence of such injury during invasive endotracheal intubation maneuvers^30^. Nevertheless, *in silico* studies are limited in addressing inflammatory cascades arising in conducting airways and have instead relied on extrapolation from hemodynamic studies with endothelial cells, limiting their clinical relevance^31^.

In the present work, we explore for the first time the hypothesis of potential immunological factors originating in the upper respiratory tract as a result of ventilatory flow-induced shear stresses. To this end, we developed a true-scale, 3D bronchial epithelial airway *in vitro* model of the tracheobronchial tree subject to physiologically-realistic ventilatory protocols in intubated pediatric populations that are prone to VILI^32^. We expose injurious effects of flow-induced WSS on the epithelial airway barrier populating the 3D airway lumen, leveraging phenotypical endpoints of epithelial structural integrity, cell apoptosis, and importantly the secretion of cytokines associated with inflammatory pathways. Our *in vitro* assays support the manifestation of shear flow-induced lung injury during mechanical ventilation, thus strenghening the biotrauma hypothesis. To mitigate the initiation of such inflammatory cascades during ventilation, we demonstrate as a proof-of-principle the topical delivery of a widely used anti-inflammatory respiratory therapeutic for preventive action as a prophylactic strategy that supports opportunities in preclinical pulmonary research towards protective therapies.

## Results

### Development of 3D bronchial epithelial airway model

As preterm infants are a unique patient population that may exhibit significant damage resulting from mechanical ventilation support^32^, we engineered an *in vitro* neonatal upper airway model (Figure 1a) to investigate the effects of flow-induced shear stresses on the tracheobronchial epithelium during intubated ventilation. The model spans the trachea to the first three bronchial generations of the airway tree (Figure 1b) and adheres^33,34^ closely to the idealized planar double-bifurcating Weibel^35^ lung model. Following previously reported methods^30,36^ we homothetically scaled the model’s geometric dimensions to the anatomical size of a ∼2 kg (33 weeks old) preterm infant based on the tracheal diameter. We used additive manufacturing (i.e., 3D printing) to fabricate a cast (Figure 1c) and filled it with liquid polydimethylsiloxane (PDMS). Following polymerization, we dissolved the printed material leaving a transparent PDMS phantom (Figure 1d) used as an anatomically-realistic scaffold architecture for cell culture. Next, extracellular membrane (ECM) proteins (i.e., fibronectin and collagen) were used to coat the model’s inner surface before cell culture following an optimized coating study (see Supplementary Figures 6,7). Here, we cultured Calu-3 cells (Figure 1e); Calu-3 is an established human bronchial epithelial cell line and widely used in preclinical pulmonary research for drug screening and toxicity^37,38^, thereby recapitulating a relevant epithelium lining the bronchial regions. Specifically, using an airway epithelial cell line permits experimental control and reproducibility as a preclinical “gold standard” benchmark by reducing variability in cell cultures arising from donor to donor differences^39^ while maintaining key features of the bronchial epithelium in human lungs.

**Figure 1:**
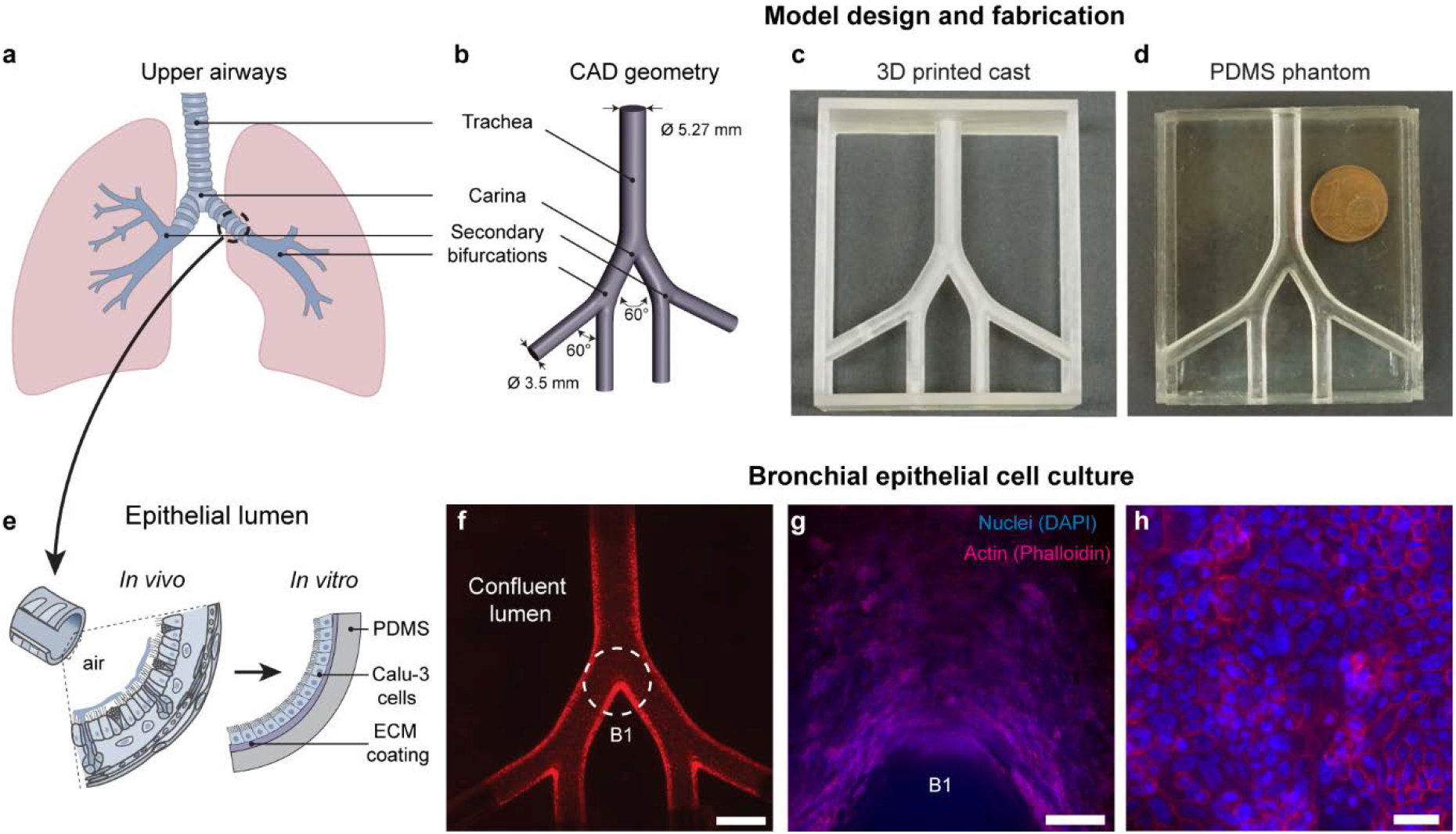
Development of a true-scale, 3D bronchial epithelial airway model. (**a**) Schematic of the lungs including the conducting upper airways. (**b**) Computer-aided design (CAD) of a simplified, symmetric, double-bifurcating model starting at the first generation of the airway tree (i.e., trachea) up until and including the third bifurcating generation of bronchi. The model’s dimensions match the scale of a 33 week old (preterm) infant. (**c**) 3D printed cast used to fabricate a (**d**) transparent, polydimethylsiloxane (PDMS) phantom photographed alongside one euro (€) cent coin for scale. (**e**) The *in vivo* epithelium-lined lumen is recapitulated *in vitro* by culturing bronchial Calu-3 cells on the PDMS surface coated with extracellular matrix (ECM) materials (i.e., fibronectin and collagen). (**f**) The model is imaged via stereomicroscope and red fluorescent membranal staining (CellTracker™ Red CMTPX), exhibiting a fully confluent monolayer of cells covering the inner lumen surface. Scale bar in (f) is 5 mm. A dashed white circle highlights the first bifurcation (tracheal carina), shown at two magnifications: (**g**) 4x and (**h**) 20x after staining F-actin (Phalloidin-iFluor 555) and nuclei (DAPI). Scale bars for (g) and (h) are 200 µm and 40 µm, respectively. See Methods for more details on fabrication and cell culture.

Models were cultured under immersed conditions and tracked over 3-4 weeks until a fully confluent epithelial monolayer populated the entire 3D airway lumen (Figure 1f). As recently shown *in silico*, jet flow impaction in airways can be significant during invasive mechanical ventilation^40^. Hence, monitoring cell structural integrity is critical (see tight junction occludin protein staining in Supplementary Figure 7), in particular at the bifurcations (e.g. main carina in Figure 1f-h). Our development of more realistic 3D *in vitro* airway morphologies is supported by recent studies where Calu-3 cells (as well as other epithelial cells) cultured on curved membranes and inside lumen exhibit distinct characteristics from traditional 2D monolayers, including cell density and shape, apoptotic ratios, and cross-sectional morphology^41,42^. Here, we specifically designed a planar bifurcating geometry to facilitate microscopy imaging by limiting the model’s vertical dimension, whose 3D curvature of the inner lumen extends outside the focal depth under higher magnifications (Figure 1h). The total volume enclosed within the model is 1.5 ml, efficiently and robustly removed or exchanged via plastic connector ports inserted into all outlets for model maintenance, allowing for the simple removal of medium and cell collection for further analysis such as ELISA and flow cytometry (see below).

### *In vitro* and *in silico* flow dynamics reveal focal shear stresses under invasive mechanical ventilation

The intricate flow patterns in the human respiratory airways are mainly driven by the interaction of time-dependent flow fields in the upper airway generations (starting in the trachea) with changing geometries, including changes in cross-sectional areas, wall curvatures and carinal edges across the airway tree^33^. We studied both in experiments and simulations the flow dynamics in our model by imposing symmetric, sinusoidal flowrate profiles (Figure 2a) at the model’s inlet mimicking the active inhalation and exhalation phases under intubated ventilation conditions (see Table 1). We experimentally measured ventilation flows in a nearly identical PDMS phantom model (Supplementary Figure 10) to specifically improve optical access (with the addition of transparent side windows) and robustness (i.e., nylon tube connector in place of the endotracheal tube of comparable inner diameter). Transient, temporally-resolved 3D velocity vector fields were measured via tomographic particle image velocimetry^40^ (TPIV) (Figure 2b-c). For the TPIV set-up (see Supplementary Figure 11), four cameras at specifically offset angles were used to simultaneously image the flow in the model, fully capturing 3D flow features. Further processing involves triangulating the location of flow-tracing particles in a volume between the multiple cameras to resolve the flow in 3D space (see Methods). Analysis of the extracted flow fields reveals that a flow jet exiting the endotracheal tube and impacting on the main carina is the most dominant flow feature in the intubated upper airways, in particular at peak strength midway during the inhalation phase (Figure 2a), and supporting recent findings in a study using a six-generation intubated neonatal upper airway model^30^.

**Table 1:** Mechanical ventilator settings.

|  |  |  |
| --- | --- | --- |
| Positive End-Expiratory Pressure (PEEP) | 7-8 | cmH<sub>2</sub>O |
| Peak Inspiratory Pressure (PIP) | 25 | cmH<sub>2</sub>O |
| Respiratory rate | 60 | bpm |
| Inspiratory time | 0.35 | s |

**Figure 2:**
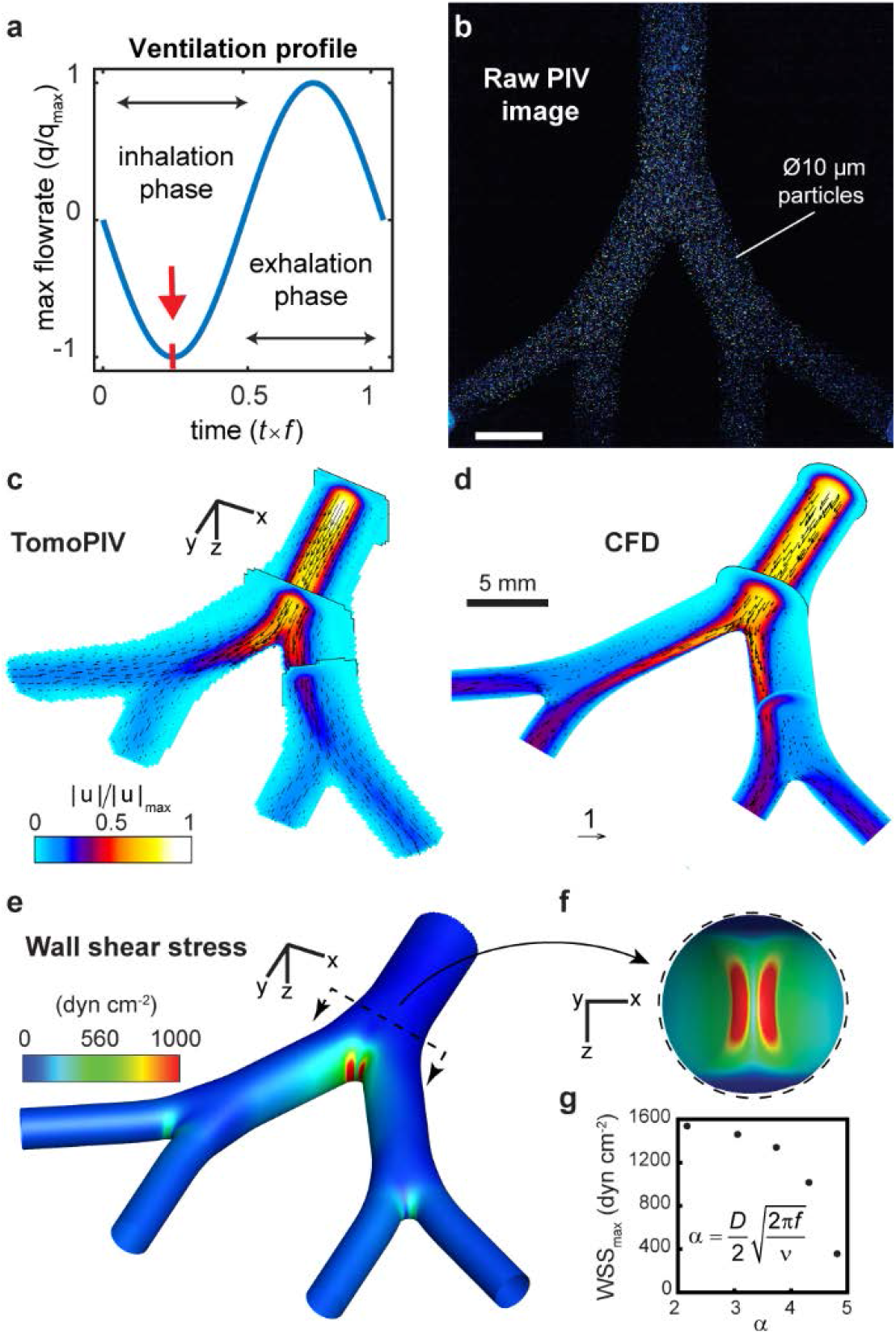
*In vitro* and *in silico* fluid dynamics analysis reveals a region of elevated shear stress concentrated at the tracheal carina during the inspiratory phase of intubated ventilation. (**a**) A sinusoidal flow profile mimicking oscillatory mechanical ventilation is imposed on the model via perfusion at the inlet (trachea). The first quarter of the cycle (see red arrow) marks the inspiratory phase’s peak strength when air is pushed into the model. The flow is visualized and measured experimentally using tomographic particle image velocimetry (TPIV), demonstrated with a raw image (**b**) from one of four cameras in the TPIV set-up (see Methods and Supplementary Figures 8, 9), showing illumination of 10 µm diameter fluorescent particles captured instantaneously while tracing the streamlines of the flow. Following image analysis (i.e., image pre-processing and TPIV algorithms), the 3D transient flow is fully resolved. The flow field at peak inspiration is plotted in (**c**), with several orthogonal cut planes colored by the normalized velocity magnitude contour field and overlayed with velocity vectors. (**d**) An *in silico*, i.e., computational fluid dynamics (CFD), solution is compared with the experimental data. During peak inspiration, the most dominant flow feature is captured by a synthetic jet exiting the endotracheal tube and impacting the first carina. A numerical solution allows for finer near-wall resolution and analysis of wall shear stresses (WSS), mapped by colored contours in (**e**) with a top view of the first bifurcation shown in the inset (**f**). Panels **b-f** feature a flow analysis of a representative low-frequency ventilation protocol most similar to conventional mechanical ventilation used in cellular *in vitro* experiments. In (**g**), maximum wall-shear stress (WSS) levels measured at the first bifurcation are plotted as a function of normalized ventilation frequency, or *α* (i.e. Womersley number), for five ventilation protocols solved using CFD (see Supplementary Figure 11).

We next ran *in silico* computational fluid dynamics (CFD) simulations modeling identical flow conditions in the airway model over several ventilation flow profiles using the finite volume method (FVM) and compared results with experimental measurements for validation. In Figure 2d, we plot a numerical solution matching the experimental case conditions shown in Figure 2c, with orthogonal cut planes colored according to the normalized velocity magnitudes and an overlay of velocity vectors. Given the spatial imaging limitations in TPIV experiments^40^, CFD is particularly useful in resolving flows at higher spatial resolution, in particular at the carina and in the vicinity of the lumen walls. In Figure 2e, we plot the computed wall shear stresses (WSS) on the model lumen during peak inspiration when the inhalation jet is at peak strength. We find an elevated WSS region localized at the jet’s impaction site with the main carina, shown in an enlarged inner side view in Figure 2f. Here, local WSS values exceed 1,500 dyn·cm^-2^; namely two orders of magnitude higher than levels found to impair epithelial permeability in human and mice bronchial epithelial cells^43^. We find lower (<300 dyn·cm^-2^) WSS values concentrated in the two daughter bifurcations, an ∼80% attenuation relative to the first generation due to viscous energy dissipation and anticipated to occur in any symmetrically branching channel system^44^. We performed four additional simulations with smaller tidal volumes but identical flow rates (i.e. breathing frequency is increased according to the linear relationship^30^ *Q* = *f*×*TV*) and tracked the peak WSS values at the main carina. In Figure 2g, results are plotted as a function of the Womersley number, defined as 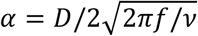 a non-dimensional frequency term varied between 2 and 5. A reduction in maximum WSS values relative to *α* indicates that lower tidal volumes at higher frequencies attenuate the flow jet’s effects, implying a possible link to the acknowledged protective benefits of low tidal volume ventilation^45^.

We next compare these findings to previous numerical studies on flow-induced shear stress in human airways and find similarly reported patterns of local peaks at the bifurcating sites. In an airway model of generations 3-5, elevated wall shear stresses (∼5 dyn·cm^-2^) were reported^46^ localized at the bifurcations, matching our findings that the primary site for wall shear stress is indeed at the impaction site of the intubation jet with the main carina and that subsequent bifurcations witness the localization of far lower shear stresses. A large eddy simulation (LES) type computational study found that the addition of an endotracheal tube (ETT) in a computed tomography (CT) scan patient-derived airway model increased wall shear stresses at the main carina during inspiration by ∼87% to a peak of ∼200 dyn·cm^-2^. These values are likely lower than our peak WSS values as their study focused on high-frequency ventilation protocols (10<*α*<20) using smaller tidal volumes. Our flow analysis corresponds to an endotracheal tube positioned axisymmetrically in the trachea of our transparent model. In clinical practice, medical imaging (e.g., X-ray) is required to confirm proper ETT placement; a situation that may be altered by the infant’s movement and changes in body position. Muller *et al*.^47^ studied the effects of ETT positioning on tracheal wall shear stress under high-frequency jet ventilation (HFJV), concluding that shear forces were amplified more than twice in cases where ETT tubes were placed asymmetrically more proximal to the tracheal wall, indicating our relatively conservative approach for the ETT placement.

### Replicating clinical settings of invasive ventilation *in vitro*

Once models achieved full epithelial confluence following 3-4 weeks of culture (Figure 3a), these were prepared for exposure to invasive ventilation, replicated with equipment and corresponding clinical settings typically used in neonatal intensive care units (NICUs), as described schematically in Figure 3b-c. A 3.0 mm inner diameter (ID) uncuffed endotracheal tube was inserted 2 cm above the first bifurcation apex (see enlarged inset in Figure 3b), matching approximately the insertion depth of 8 cm for a 2 kg (∼33 weeks) infant following the American Academy of Pediatrics guidelines^32^. Clinical ventilation settings were maintained (in the absence of the lower regions of the lung) by fitting compliance adapters (small latex balloons, see Figure 3b) to the four outlets of the model and calibrated to match clinical pressure and lung compliance conditions monitored by the ventilator (see Methods). A natural rubber latex membrane was similarly used and calibrated in simulating lung compliance of distal airways *in vitro* in a study on the role of ventilation mechanics in reopening a collapsed lung^48^. An additional model was placed alongside and connected to identical tubing and compliance adapters without active flow ventilation serving as a control for the ventilation exposure test.

**Figure 3:**
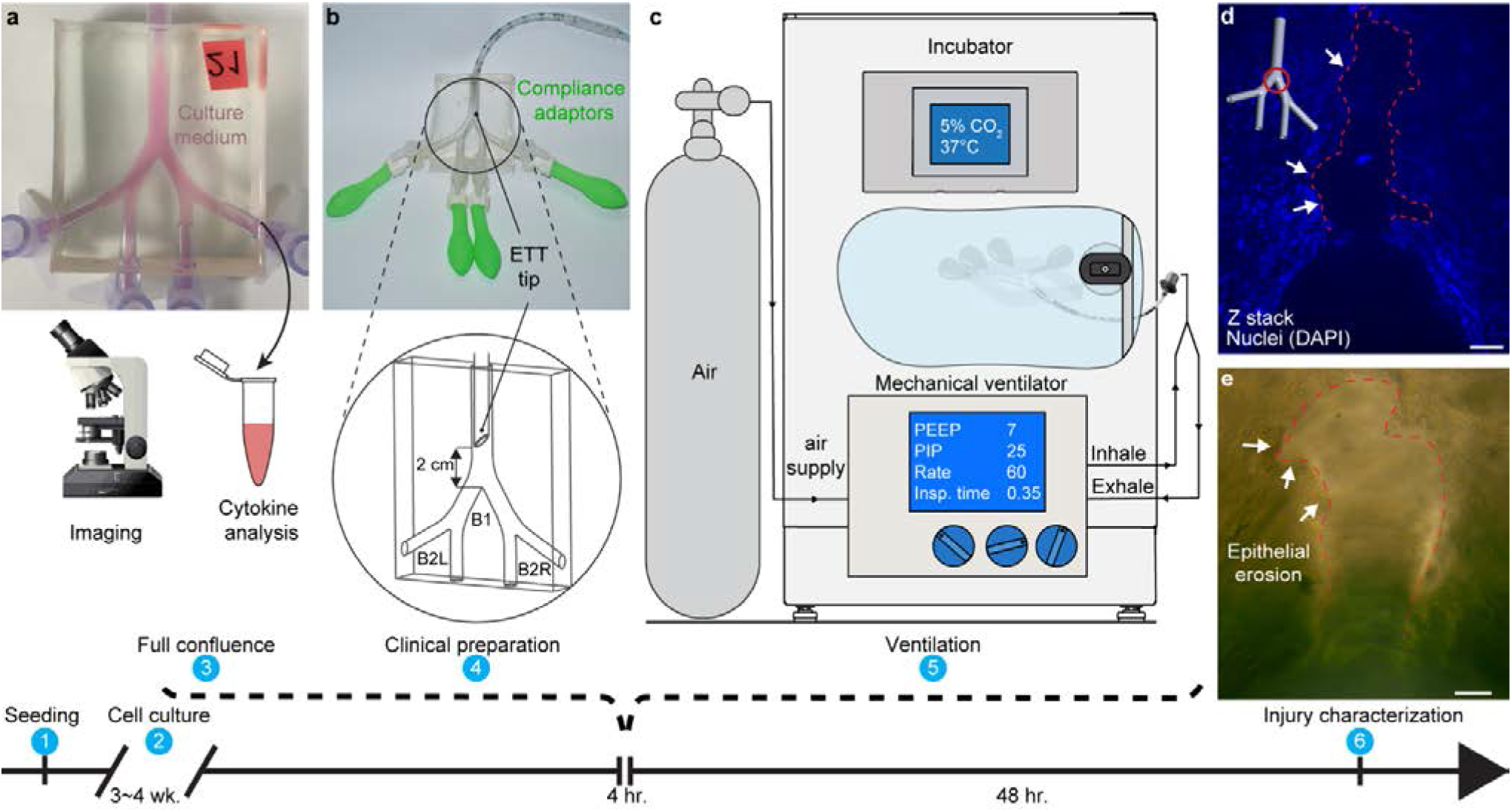
Simulated mechanical ventilation experimental set-up and detection of cellular injury. (**a**) The 3D lumen epithelium is recapitulated by seeding (1) bronchial cells (Calu-3) inside the model and cultured (2) at 37 °C and 5% CO_2_ until full confluence (3) is achieved (3-4 weeks). Before exposure to either ventilation or control conditions, models are imaged, and their culture medium collected for later cytokines analysis, serving as a reference point for a priori conditions. (**b**) Next, the model is prepared (4) for clinical ventilation by installing compliance adapters (latex balloons) at the four outlets (to simulate pressure conditions present in a full lung) and inserting a 3.0 mm ID uncuffed endotracheal tube (ETT) 2 cm above the first carina (marked B1), following clinical guidelines for proper insertion depth. The two daughter branches are marked B2L (left) and B2R (right) following anatomical orientation convention. Note that during ventilation, the ETT tip is placed symmetrically while here shown at a 90 deg. rotation for clarity. (**c**) Models are ventilated (5) inside the incubator with medical-grade air, supplied via a breathing circuit connected to a neonatal mechanical ventilator for 4 h using settings defined in Table 1. After the exposure, models are filled with medium and returned to the incubator for an additional 24-48 h before further analysis and injury characterization (6), including microscopy imaging. (**d**) Fluorescent and (**e**) bright-field microscopy imaging reveals a region of cell detachment localized at the first bifurcation, exhibited 48 hours following invasive ventilation. The delayed onset of epithelial erosion, i.e., a severe form of cell injury, demonstrates the complex traumatic pathway following exposure to flow-induced shear stresses during invasive ventilation. Regions of cell detachment are highlighted with thin dashed lines. (d) Nuclei stained with blue DAPI. Scale bars in (d) and (e) are 100 µm.

To explore ventilation conditions on the epithelium at the air-liquid interface (ALI), models were emptied of culture medium (stored for cytokine analysis, see below), connected to ventilation tubing, and ventilated for 4h. Models were kept inside an incubator (37 °C, 5% CO_2_) for the duration of the exposure to factor for environmental stress not associated with the ventilation. Ventilator settings (Table 1) were chosen on the higher side of the recommended clinical range, used for infants with significant breathing difficulty and at a low respiratory rate^49,50^ (i.e., frequency). These settings were selected to underline the potential for damage, following our flow analysis revealing the effects of shear stresses under intubated ventilation conditions, and highest at lower respiratory frequencies. Following the 4 h exposure assay, models were refilled with fresh cultured medium and returned to the incubator. Later this medium was collected at identical culture time intervals as pre-exposure samples.

In line with our experimental flow analysis identifying localized, elevated wall shear stress sites (Figure 2), we found evidence of physical damage to the epithelium in our model at the same locations following exposure to intubated ventilation conditions. Epithelial erosion (i.e., the detachment of epithelial cells from the model) was observed using fluorescent nuclei staining (Figure 3d) and bright-field microscopy (Figure 3e) at the first bifurcation (i.e., main carina) 48 h following a simulated ventilation exposure. We tracked the evolution of epithelial erosion by imaging in bright-field the live cells in the models at several time points following ventilation and compared it to the initial pre-ventilation image at the same locations (see Supplementary Figure 8). Erosion was not observed in other regions of the ventilated model, including the two daughter bifurcations where minor shear stress localization was predicted by the flow analysis (Figure 2e). Notably, epithelial erosion in the tracheobronchial region is known as a severe histopathological marker of inflammation and acute lung injury found in pathological slides sampled from infants who died following mechanical ventilation^51,52^. The prevalence of this damage in invasively ventilated infants today is mostly unknown due to significantly lower mortality rates and autopsy protocols that may not include specific microscopic examinations of the trachea, carina, or mainstem bronchi. Moreover, epithelial restoration mechanisms^53^ may repair denuded regions in surviving infants, further obscuring potential observation of this phenomena in the clinic. Solid mechanical stresses in the context of cyclic reopening were shown to contribute to alveolar epithelial cell death and detachment in a microfluidic model^54^ and may similarly be a suitable proxy here for identifying structural lung pathology *in vitro*.

### An increase in cell apoptosis indicates an early *biotrauma* cascade

Following ventilation/control exposure experiments, cells were harvested from the models and analyzed for cell death using fluorescence-activated cell sorting (FACS) with apoptosis/necrosis using Annexin-V/ propidium iodide (PI) staining (Figure 4a-c). Annexin-V provides a sensitive method for detecting cellular apoptosis, while propidium iodide (PI) is used to detect necrotic or late apoptotic cells. Nearly all cells from the control models were found to be viable cells (double-negative) when analyzed at both 24 h and 48 h time points following exposure (Figure 4a), while large decreases in % live cells were found among the ventilated models. The %population of live cells (indicated double-negative for both Annexin-V and PI) in ventilated models decreased by 60% with 24 h following the exposure and 75% at 48 h, respecitvely. Most non-viable cells were found in the apoptotic sub-population (indicated double-positive for both Annexin-V and PI), increasing from <5% of the total cell population to 55% and 60% at 24 h and 48 h, respectively. Necrotic cells (indicated positive for PI and negative for Annexin-V) were found in negligible levels in both control and ventilated models at both time intervals, as seen in Figure 4c. In interpreting such results, we recall that cell apoptosis is a tightly regulated mechanism routinely employed in maintaining normal tissue and organ homeostasis. Macrophages remove apoptic cells via phagocytosis, a process enhanced via the local release of anti-inflammatory cyotkines interleukin IL-10 and TGF-β1 by both macrophages and apoptotic cells^55^. Incidence rates of apoptosis among airway epithelial cells above normal levels can indicate a compensatory response to disease or a pathogenetic consequence, or both^56^. Pulmonary inflammation and fibrosis are known to be linked to the accumulation of excessive apoptotic cells or the cleavage of cell remnants, which in turn may promote further inflammatory processes^55^.

**Figure 4:**
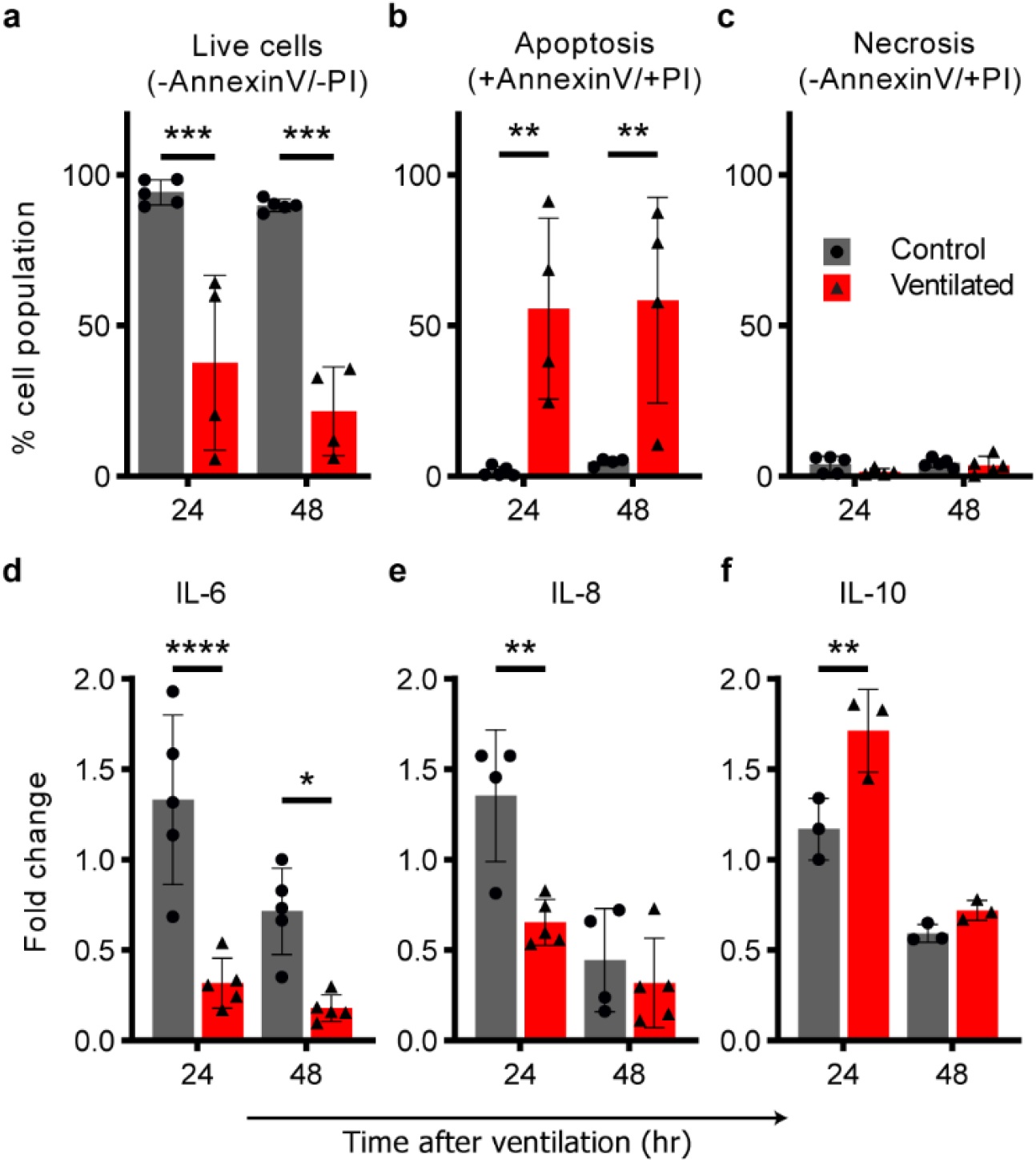
Ventilation exposure modulates apoptotic pathways and epithelial inflammatory cytokine secretions. Cells and culture medium were collected from models 24 h and 48 h following ventilation exposure and compared with baseline levels. Air ventilation was performed for 4 h, or the ventilator machine kept off for the same duration as a control. Following the exposure, culture medium was filled back into the models. (**a-c**) An increase in apoptotic cells indicates a *biotraumatic* response. Cells were collected from models and analyzed using flow cytometry (FACS) following Annexin-V and Propidium Iodide (PI) staining. The %total population of (**a**) live cells decreased by 60% at 24 h and 75% at 48 h in the ventilated group relative to the control, which is largely unaffected at either time point. Most of this increase in non-viable cells is found in (**b**) the apoptotic, ventilated group while (**c**) negligible necrotic cells were counted for either exposure group. The increase in apoptosis was consistent in models analyzed both 24 h and 48 h following the exposure experiment. (**d-f**) Changes in the secretion of inflammatory cytokines following ventilation suggest a signaling pathway. Cytokines IL-6 (**d**), IL-8 (**e**) and IL-10 (**f**) were measured in culture medium collected from the models after 24 h and 48 h periods before and following simulated ventilation, respectively. Measurements from repeated experiments are plotted as normalized fold change to highlight signaling changes resulting from the test exposure. In (**d**) and (**e**), we measured a significant reduction in pro-inflammatory IL-6 (75% at both 24 h and 48 h) and IL-8 (50% at 24 h) relative to the control, while conversely in (**f**), we find that IL-10, an anti-inflammatory cytokine, is elevated >30% at the 24 h time point compared with the control. All values are normalized relative to their baseline, i.e., divided by pre-exposure secretory levels. Cytokines were measured using ELISA. All graphs show mean (SD) values. ^*^ p<0.05; ^**^ p<0.01; ^***^ p<0.001; ^****^ p<0.0001. Source data are provided as a Source Data file.

### Ventilation exposure modulates epithelial inflammatory cytokine secretions

To explore the causal relationship between biophysical injury and induced inflammatory response, the central tenet to the *biotrauma* hypothesis^2^, we tracked the secretion of cytokines known to be critical mediators of pulmonary injury and inflammation. Secreted concentrations of pro-inflammatory (IL-6, IL-8) and suppressory anti-inflammatory (IL-10) cytokines were measured in the sampled cultured medium at two intervals: 24 h and 48 h before and after the ventilation/control exposure. In Figure 4d-f, cytokine secretions are plotted as fold change relative to baseline, pre-exposure levels. We found decreased IL-6 and IL-8 among the ventilated models at both time intervals, while the control group showed slightly increased levels. At 24 h and 48 h, IL-6 decreased by more than half of baseline levels and ∼75% relative to the control; IL-8 decreased ∼25% at 24 h and 50% relative to the control group, with no statistical significant reduction at 48 h. Conversely, elevated concentrations of IL-10 were measured 24 h after ventilation while the control remained close to baseline levels. All cytokine secretory events were observed to be most prominent in the first 24 h following the exposure, while at 48 h effects were found diminished, indicating the transitory nature of the response to a single, 4 h ventilation exposure protocol. Notably, IL-10 is known to play a crucial role in regulating immune responses by limiting and ultimately terminating inflammatory events via the inhibition of cytokine synthesis^57^. One possible explanation for our findings is that at 24 h post-stimulus, pro-inflammatory cytokine production is inhibited by IL-10 due to an anti-inflammatory response.

### Prophylactic leukotriene receptor antagonist reduces cell death and modulates the secretion of inflammatory cytokines

After establishing the presence of an inflammatory response to flow-induced shear stress in our model (Figure 4), we next sought a direct modulatory agent that could demonstrate the model’s application for therapeutic preclinical research. As a proof-of-concept, we tested the prophylactic effect of the commonly used airway anti-inflammatory drug Montelukast (sold under the brand name Singulair^®^) of the leukotriene receptor antagonist (LTRA) family. LTRAs have bronchodilatory and anti-inflammatory effects, typically used together with inhaled corticosteroids (ICS) in the maintenance treatment of adult and pediatric asthma^58^. We injected culture medium supplemented with Montelukast (0.006 µg/ml) into models for 2 h prior to cellular exposure at the ALI to either ventilation or control conditions. The models were subsequently analyzed for cell death (i.e., apoptosis/necrosis) and cytokine secretions (i.e., IL-6, IL-8 and IL-10) following the previous section’s protocol. Figure 5a-c plots cell cytometry analysis following apoptosis/necrosis staining 24 h following the ventilation/control exposure, identified previously as a critical time point in developing a biotraumatic response. In Figure 5a, we report a significant increase (55%) in the total percentage of live cells analyzed from ventilated models treated prophylactically with Montelukast relative to models ventilated with no prior treatment. The increase of live cells in the prophylactically treated, ventilated group follows the reduction (75%) in apoptotic cells (Figure 5b) in this same group. Though the percent population of necrotic cells in all groups was small (1-5%), as shown in Figure 5c, we measured a significant increase (70%) in necrotic cells in the Montelukast treated, ventilated group. We found no significant change due to prophylactic treatment in either control group, with viable cell count measured >95% in all non-ventilated models.

**Figure 5:**
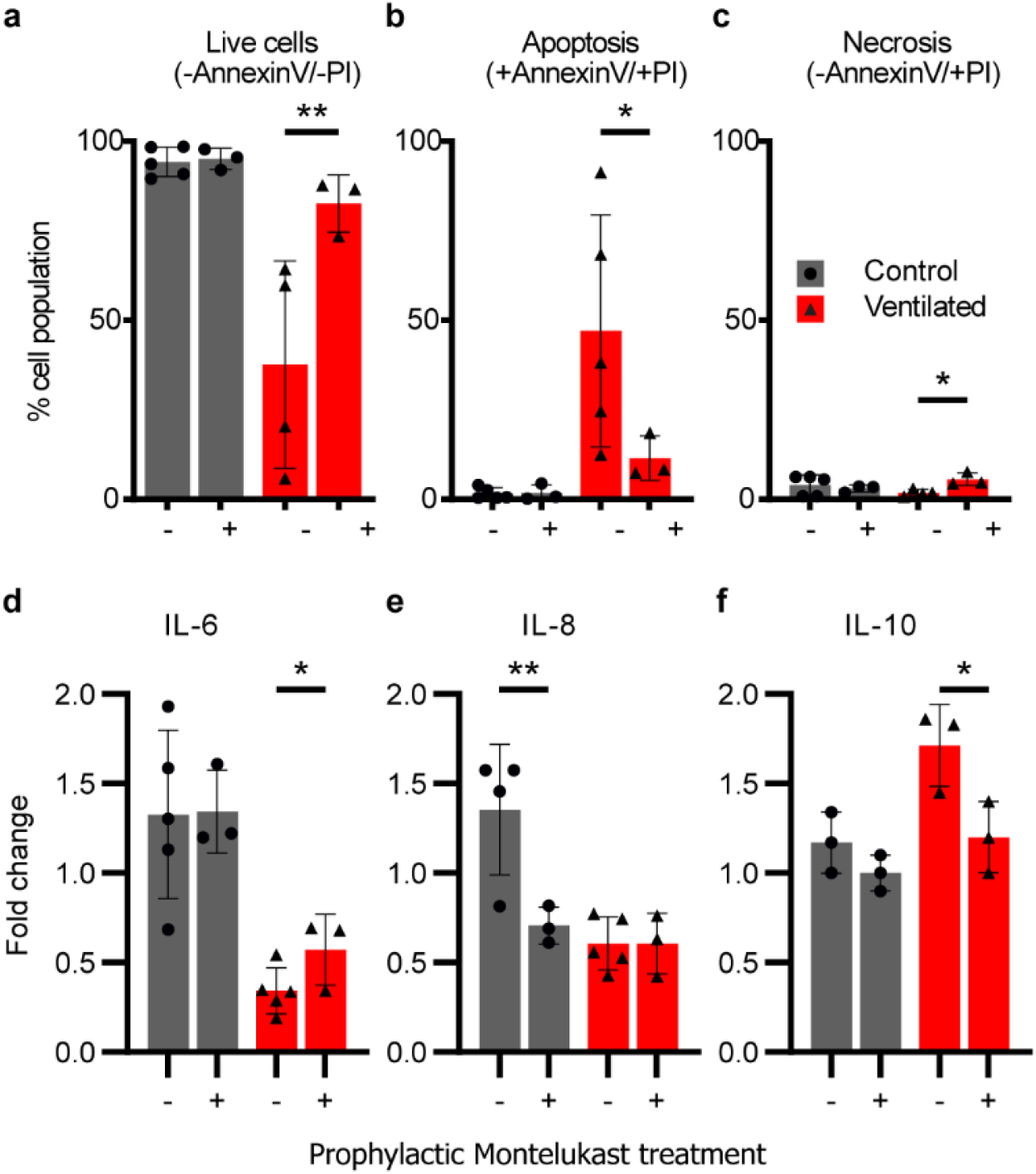
A prophylactic leukotriene receptor antagonist (Montelukast) reduces cell apoptosis and modulates inflammatory cytokine secretion in bronchial epithelial airway models exposed to ventilation injury. Montelukast, an asthma anti-inflammatory medication commonly used to treat pulmonary asthma, was supplemented in the culture medium of models for 2 h before ventilation exposure. Flow cytometry was performed on cells collected from models 24 h following the ventilation/control and stained with Annexin-V and Propidium Iodide (PI), indicating % of the cell population (**a**) live, (**b**) apoptotic (**c**), or necrotic. Montelukast pre-treatment significantly increased (55%) the number of live cells in the ventilated group by reducing apoptosis (75%) while showing no adverse impact on the control group’s high live cell count. (**c**) An increase in necrotic cells (70%) was observed in the ventilated group while absent in control, with a negligible effect on %total of live cells due to low overall counts (maximum of 5% total population). (**d-f**) Cytokine secretions were measured in the culture medium collected from all models 24 hours after the ventilation exposure. Cytokines IL-6 (**d**), IL-8 (**e**), and IL-10 (**f**) are plotted as fold change normalization to highlight changes in signaling resulting from the test exposure. (**d**) We report no significant difference in IL-6 secretion in the control groups, while an increase in IL-6 secretion (67%) was measured in the pre-treated ventilated group relative to non-treated. (**e**) In contrast to IL-6, IL-8 secretion was found reduced in the control group (50%) following prophylactic treatment, while we measured no difference due to treatment in the ventilated groups. Lastly, we measured the suppressory IL-10 secretion, with mean values unaltered in the control groups but reduced by 30% following prophylactic treatment with Montelukast. All cytokine secretion values are plotted as fold change, i.e., normalized relative to their baseline pre-exposure levels. Cytokines were measured using ELISA. All graphs show mean (SD) values. ^*^ p<0.05; ^**^ p<0.01; ^***^ p<0.001; ^****^p<0.0001. Source data are provided as a Source Data file.

In Figure 5d-f, we report the effect of prophylactic Montelukast treatment on inflammatory cytokines (IL-6, IL-8 and IL-10). In the ventilated group, where we previously measured a reduction in pro-inflammatory IL-6 and IL-8 following ventilation exposure, we see a similar reduction (Figure 5a-b), though values of IL-6 are significantly higher (67%) in the pre-treated group relative to non-treated. We note that while IL-8 secretory levels were unchanged in the ventilated group following prophylactic treatment, a reduction of 50% was measured in the control (i.e., no ventilation exposure) group. For suppressory IL-10 (Figure 5f), we measure a reduction (30%) to near-baseline levels in the prophylactically treated ventilated models, a reversal relative to the non-treated exposure group. These results suggest that the inflammatory response elicited in our models by ventilation exposure is directly modulated by prophylactic treatment using Montelukast.

## Discussion

This study aimed to shed light on the physical pathways contributing to *biotrauma*; the type of ventilator-induced lung injury most poorly defined and understood^4^. Preterm infants are particularly susceptible to extended periods of ventilatory support; the more immature the newborn is at birth the longer the likely period of ventilation and thus the increased risk of respiratory distress severity^65^. Moreover, preterm infants are considered more susceptible to external stressors as their respiratory organs are not fully matured and functional^65^. Due to their prolonged exposure to ventilation and their anatomical scales, we based our *in vitro* airway model on the true-scale anatomy of preterm infants undergoing invasive ventilation. Unlike adult patients suffering from lung disease, the preterm infants’ central condition lies in their underdeveloped lungs typically exluding other morbidity, thereby isolating the effects of ventilation from the other underlying conditions such as smoking and diabetes, common among adult ventilated patients^65^.

Following a recent theoretical and computational analysis^30^, we hypothesized that flow-induced shear stresses in the tracheobronchial tree’s upper regions could be associated with a biotraumatic injury. We designed and constructed a 3D tracheobronchial airway model featuring a fully-confluent cultured epithelial cell monolayer. We exposed our model to physiologically-relevant ventilation flow conditions and used quantitative fluid mechanics tools to identify the localized airway “hot spots” most susceptible to stress-related injury under intubated mechanical ventilation. We found evidence of epithelium structural destruction (i.e., cell erosion) localized at the impaction site between a flow jet emanating from the endotracheal tube tip and the main carina.

Advanced *in vitro* platforms have introduced new bioengineering methodologies to explore human diseases *in vitro* by recapitulating key physiological functions in isolation, thereby offering new insights that are often well beyond reach in vastly more complex *in vivo* models^19^. For example, *lung-on-chips*^21,59^ have established an attractive strategy to study amongst other human lung inflammation and drug responses *in vitro* by recreating a human-relevant alveolar microenvironment. To date, some studies have investigated ventilator-induced lung injury (VILI) and modeled the downstream barotrauma effects of pressure and membranal stretching at the acinar microscales only^41,54^ whereas convective flow phenomena occurring upstream at the macroscale (i.e. conducting airways) have been largely overlooked^60^. Yet, upper airway flow phenomena drive downstream effects and thereby influence whole-organ lung function. For VILI studies, omitting the lung macroscale potentially implies overlooking the primary insult’s origin, namely where the invasive medical intervention at the root of the problem first meets the patient. A gap thus exists between macroscale physiological flow phenomena characteristic of actual clinical settings and the downstream flow conditions introduced in microscale *in vitro* models. Here, our hypothesis for flow-induced *biotrauma* is rooted in the fluid mechanics of the clinical environment and the derivative shear forces imposed on the macro architecture of the upper airways.

We sought a simple and robust *in vitro* model for exploring the intubated ventilation jet’s interaction with the upper airway’s epithelium. Extrapolating our computational data and comparing it to other studies that included a broader anatomical scope found that the flow jet’s effect is negligible beyond the first several airway generations. Therefore, we designed a model that included the trachea and first three bronchial generations, a relatively limited section of the full respiratory tract. Our flow physics analysis confirmed that the jet’s effect is mostly dissipated upon entrance into the second generation in our model due to energy losses beyond the first bifurcations. To explore the cellular response to the introduced flow phenomena, we used our reconstructed airway model’s PDMS surface as a scaffold for culturing a monolayer of bronchial epithelial cells, covering the entire three-dimensional lumen. We chose Calu-3, a well-established lung cell line for preclinical pulmonary research that grows confluent layers in immersed conditions and forms barrier junctions. Previous *lung-on-chip* models have recapitulated other essential aspects of epithelial cell function such as ciliation^61^ and mucus^41^ secretion by culturing primary cells at an air-liquid-interface, but have excluded physiologically-relevant flow conditions critical to our study. Mucus is an aqueous secretion that offers a critical protective lining to epithelial cells covering the *in vivo* airways and serves to clear contaminants upwards via ciliary motion. While intubated, however, ventilated patients are typically suctioned of secretions^62^, including mucus, to quickly clear the airways and facilitate breathing. Following clinical guidelines^62^, our ventilated model similarly excludes mucus, though its potential for protecting against the injury we have identified here warrants further research.

To isolate the effects of mechanical stress from intubated ventilation, we exposed our models to 4 h of clinical ventilation conditions in air; a duration that was short enough to avoid introducing interference owing to the removal of cells from maintenance conditions, i.e., immersion in culture medium but sufficiently long to observe a clear cellular response. Furthermore, a 4 h exposure protocol follows previous studies investigating intubated ventilation in “one hit” *in vivo*^15,63^ and *in vitro* models^64^, facilitating the interpretation of our results relative to the broader literature. In the clinical setting, patients can be supported by a ventilator for days or even weeks, likely amplifying the effects measured in our *in vitro* models following exposure to 4 h alone.

In analyzing our models following exposure to ventilated conditions, we first explored changes to the epithelium layer’s structural integrity. We found evidence of cell denudation localized primarily at the main carina, matching our flow analysis that identified concentrated levels of high shear stress at the same site, at maximum during the inhalation phase of simulated ventilation. The deleterious effects of a jet stream in the tracheobronchial region were first suspected in the context of high-frequency jet ventilation (HFJV); a strategy of delivering short pulses of pressurized gas directly into the upper airway through a custom-designed endotracheal lumen. Proposed initially to improve gas exchange and thereby reduce the severity of VILI in both infants^66^ and adults^67^, HFJV is no longer recommended for use following a meta-analysis that failed to find an advantage in reducing mortality rates^68^, in addition to clinical^51^ and animal^69^ studies revealing acute adverse effects (e.g. necrotizing bronchitis, epithelial erosion, loss of surface cilia, and other markers of inflammation). Limitations in trial design and imprecision due to the small number of infants available for these clinical studies increases the possibility that the damaging effects of a jet stream during mechanical ventilation is not restricted to HFJV but could instead apply to all intubated ventilation protocols and not fully appreciated until now.

Previous studies may have also overlooked the potentially harmful effect of intubated flows due to differences in anatomy between humans and laboratory animal species. Rodents have monopodial airway branching, in contrast to the regular dichotomous branching in humans^70^. This situation likely alters the wall shear stress distribution from intubated ventilation though the effects of these anatomical variations on airflow and have not explicitly been explored. Even within humans, differences in anatomy and airflows exist between pediatric and adult populations, a topic of renewed interest, particularly in pharmacological dosing of respiratory drugs^71^. Here, we found that epithelial erosion exhibited 48 h after the 4 h ventilation exposure in our simplified, symmetrically bifurcating airway anatomy. While a lengthier exposure may have incurred more extensive damage, the delayed occurence of cell structural destruction indicates that the ventilation jet’s most immediate impact within such timeframe is likely to be a more subtle and complex biological response rather than a singular, acute phenomenon.

To further explore the cellular response in our model and the possible development of biotrauma following invasive ventilation, we measured cell death (apoptosis/necrosis assay) and cytokine secretions. Models were analyzed at 24 h and 48 h following exposure, with viability in the ventilated group reduced by 60%, with more than half to apoptosis, while the controls retained over 95% live cells. Measurement of cytokines secreted in the culture medium revealed increased secretion of an anti-inflammatory mediator (IL-10) while simultaneously measuring reduced secretions of pro-inflammatory IL-6 and IL-8 linked to ventilation exposure^45,57^. Owing to its anti-inflammatory effects, IL-10 has been proposed as a potential therapeutic after showing protective benefit in mice given nebulized IL-10 before injurious ventilation^15^. Interpreting our cytokine measurements could indicate the beginning of a healing process^53^ within our model following a 4 h exposure period, though the role of mediators involved in respiratory inflammation and VILI remains controversial owing to inconsistent results^15^. In a rat model of acute respiratory distress syndrome (ARDS), Chiumello *et al*.^14^ showed that local and systemic pro-inflammatory cytokines increase over time after the beginning of injurious ventilation, with a high peak between 2 h and 4 h. In a human randomized control trial, however, concentrations of IL-6 and IL-8 in bronchoalveolar lavage fluid and blood serum samples from ventilated ARDS patients showed differences 36-40 h following ventilation, whereas levels remained close to baseline when measured between 24 h and 30 h after ventilation. While we found that changes in cytokines were most significant at 24 h following ventilation, it is difficult to extrapolate our results directly to a clinical insight in either infant or adult patients. However, the mechanical damage exhibited only 48 h after the ventilation exposure suggests a much more subtle form of biological damage, tying into the hypothesis of *biotrauma* related to ventilator-induced lung injury.

To establish a proof-of-concept for our model’s application as a preclinical *in vitro* platform for drug screening, we tested our system’s response to Montelukast (Singulair^®^), an anti-inflammatory leukotriene receptor antagonist (LTRA) commonly used via oral formulation in treating or preventing bronchial asthma among pediatric and adult patients^58^. In preterm infants, corticosteroids are still the mainstay for immunosuppression, proven to improve short-term lung function. However, newly uncovered adverse effects on neurological development^72,73^ have led to the search for steroid-sparing solutions to treat inflammation in this vulnerable population. Direct topical drug delivery has the advantage of significant efficacy at the anatomical site of interest while avoiding potential drug-related adverse events which may develop following the systemic administration of formulations. Montelukast’s potential for reducing inflammation among premature infants would carry significant clinical value. The prophylactic topical delivery of Montelukast in our models demonstrated that the cellular and inflammatory response induced by exposure to clinical ventilation conditions could be modulated pharmacologically. In models pre-treated with Montelukast and subsequently exposed to injurious ventilation, we found increased cell viability together with changes to cytokine secretions, aligning with clinical outcomes found *in vivo*^74–76^. While further experiments are necessary to investigate the longer-term effects of this intervention, our findings demonstrate the *in vitro* platform’s potential as a lung biotrauma model and a potent preclinical tool for identifying new therapeutic opportunities.

## Methods

### Model fabrication

Fabrication of the neonatal tracheobronchial model (Figure 1a) begins with the computer-aided design (CAD) of a planar, symmetric double-bifurcating geometry with a constant bifurcating angle of 60, as schematically shown in Figure 1b. A stereo-lithography type 3D printer (Form3, Formlabs, USA) was used to create a casting template (Figure 1c) in the CAD model’s shape shown in Figure 1b. Printing with a layer thickness of 25 μμm ensured a smooth surface and fully resolved details, yielding a total printing time of ∼24 hours. Following post-printing preparation (removal of supports and excess print material), liquid-form polydimethylsiloxane (Sylgard184, Dow Corning, UK) was poured into the cast following the manufacturer’s instructions and left to polymerize at room temperature for 24 hours. The 3D-printed cast was subsequently dissolved by immersion in acetone for ∼24 hours, leaving a transparent hollow phantom (Figure 1d). More details can be found in a previous study using similar silicone phantom fabrication methods^30^. Following the complete dissolution of the last remaining printed material, the PDMS scaffold was left to dry in a well-ventilated space (i.e., chemical hood) for >72 hours. We found that acetone’s inadequate evaporation due to shorter drying periods disrupted later cell growth and prevented full monolayer confluence.

### Model coating

Twenty-four hours before cell seeding, the inner surface of the PDMS phantom models was coated with human plasma fibronectin (#354008, Corning, NY, USA) and collagen from bovine skin (#C4243, Sigma-Aldrich, Germany) following a coating study for optimal growing conditions of Calu-3 directly on PDMS surfaces (see Supplementary Figure 6). To coax cells to grow directly on PDMS, we conducted a coating comparison study between cells growing in a 24-well plate. We took bright-field microscopy images of the wells 3-, 5-, and 7-days following cell seeding on three different coatings: 10% fetal bovine serum (FBS), 1% v/v collagen, and a combination of 1% v/v collagen, 1% v/v fibronectin in addition to no coating. The combined fibronectin and collagen coating was found to support full confluence consistently following seven days and was subsequently used in all models.

### Cell culture and seeding

Human airway epithelial cell line Calu-3 (American Type Culture Collection, Manassas, VA) was used in this study (passage number 15-20). Cells were maintained in minimum essential medium-eagle (#01-025-1B, Biological Industries, Beit Haemek, Israel) supplemented with 10% fetal bovine serum (FBS), L-Glutamine (#41-218-100, Biological Industries), and 1% antibiotic antimycotic (Sigma-Aldrich, A5955). Calu-3 cells were seeded (i.e. 10^6^ cells) in 180 cm^2^ flasks under immersed conditions and reached 80% confluency after 10 days under standard culture conditions (37 °C, 5% CO_2_ - 95% air). Next, 3-4×10^6^ cells (in 250 µl medium) were seeded inside the model. Models were then attached to a rotator (Intelli-Mixer RM-2, ELMI Ltd., Russia) overnight at 1 RPM and subsequently left under immersed conditions and reached 100% confluency after 3-4 weeks. In total, each *in vitro* model requires approximately one month to transform from a 3D printed cast to a fully confluent airway lumen. Mycoplasma controls were performed routinely and never showed infection.

### Simulated ventilation experimental set-up

An infant-pediatric ventilator (CrossVent2, Bio-Med Devices, USA) was used to ventilate the *in vitro* model with medical-grade air via a standard neonatal breathing circuit and a 3.0 mm ID endotracheal tube (#70-100-111 Blue Line Oral Nasal Endotracheal tube, Smiths Medical, UK). Simulating the function of a compliant lung in the absence of all ∼23 generations of airways was achieved by attaching tiny biocompatible water balloons (#FBA_WB-100, Wet Products, Inc., CA, USA) to the four outlets of the model and secured in place using a plastic adapter (#FCFM-001, Nordson Medical) and thread seal tape. The ventilator maintains a set pressure by adjusting the flow in a closed control feedback loop. Built-in alarm systems on the ventilator and incubator ensured consistent ventilation and environmental conditions for the ventilation exposure duration.

### Tomographic particle image velocimetry

To perform particle image velocimetry (PIV), one of the most critical requirements is that tracer particles faithfully follow the streamlines of the flow. This requirement is most often met by calculating the Stokes number, which relates the settling time of the particle to the characteristic time of the flow and ensuring it is sufficiently small, i.e., Stk≪1. To seed air and most other gases, this would require particles sizes too small to be adequately imaged in the context of our model. As typical to fluid dynamics flow visualization studies, we used instead a glycerol/water solution (58% glycerol by mass) as the working fluid affording increased density and viscosity, thereby allowing 10 µm diameter particles (PSFluoRed; microParticles GmbH, Germany) for clear optical imaging and fluorescent reflection. Conversion of experimentally measured values to their air equivalent is straightforwardly done following dimensional analysis, a standard tool in fluid mechanics to relate flow phenomena between different length scales and material properties. In facilitating the visualization experiment and preventing leaks, the endotracheal tube was replaced with a nylon tube-to-tube connector (MLSL035-1, Nordson Medical, USA) with similar dimensions but with added lips for securing tubing and placement in the model.

### Numerical methods

The system of governing equations (i.e., conservation of mass and momentum) is solved numerically using a commercial software’s finite-volume method (ANSYS Fluent v19.2). The momentum equations are discretized using the second-order upwind scheme for velocity and second-order scheme for pressure, whereas, for coupling the velocity and pressure fields, the Coupled algorithm is applied along with a least-squares-based scheme for gradients. For numerical modeling purposes, the mouth opening is treated as a velocity inlet, and two bifurcation exits as pressure outlets. Cyclic flow conditions following a sinusoidal velocity profile were applied at the inlet as defined in Figure 2a. The double-bifurcating airway model, including the intubation tube, was meshed with tetrahedral cells using a commercial meshing software (ANSYS ICEM). Rigorous mesh convergence tests were first performed to select the optimal numerical set-up ranging from 2.8M to 7.4M. The final mesh selected for all the numerical simulations is of 4.3M tetrahedral mesh converted to a polyhedral mesh with 0.74M cells approximately in Fluent (see Supplementary Figure 11a). The time step chosen for each scenario was T/500, adequately capturing the velocities (Supplementary Figure 11b-c) present in the range of the simulated ventilation profiles. Note that in matching the TPIV experiments, working fluid of glycerol/water at 58:42 ratio was used for defining the material density and viscosity at room temperature (25 °C).

### Biochemical and cytological analyses

Cytokine levels in sampled medium supernatants were measured using commercially available ELISA kits in a blinded fashion and by following the manufacturer’s directions (Invitrogen, USA) for the following cytokines: IL-6 (#88-7066-77), IL-8 (#88-8086-77), and IL-10 (#88-7106-88). For flow cytometry, cells were dissociated from the lumen of the model by adding Trypsin EDTA Solution B (0.25%), EDTA (0.05%) (Biological Industries). Following collection, culture medium (see culture) was added at a 1:1 ratio to block the Trypsin-B effect. Cells were subsequently centrifuged at 3.0 rpm for 3 min., the supernatant discarded and then resuspended in 500 µl of binding buffer, stained with 10 µl of AnnexinV-FITC and propidium iodide (PI) following manufacturer’s instructions (Annexin-V Apoptosis Staining Detection Kit, ab14084, Abcam, USA). Metabolic activity tests were performed using an alamarBlue cell viability assay (BUF012, Bio-rad, USA). The absorbance of the incubated media containing 10% alamarBlue (2hr) was measured at 570 nm and 600 nm using a microplate reader (Synergy H1, BioTek, USA).

### Prophylactic medication preparation

Montelukast sodium hydrate (# SML0101, Sigma-Aldrich, Germany) was diluted in DMSO to a stock concentration of 0.006 mg/ml and stored in -20 °C. Before a prophylactic ventilation experiment, the stock working concentration was further diluted in culture medium (MEM-eagle) at 1:1000 before injection into the models (0.006 µg/ml).

## Statistical analyses

Data are presented as the mean with error bars showing the standard deviation (SD). Statistical analyses were performed using Prism 8.0 GraphPad software (GraphPad, La Jolla, CA, USA). Data were analyzed using a un/paired two-sided Student’s t-test, One-Way ANOVA with Holm-Sidak’s post-test, Two-Way ANOVA with Sidak’s multiple comparisons test, or as indicated in Figure legends. p-values <0.05 were considered statistically significant and are reported in figures using the following notation: ^*^p<0.05; ^**^p<0.01; ^***^p<0.001; ^****^p<0.0001 (with ^*^ referring to the comparisons specified in each Figure legend).

## Data availability

All data supporting this study’s findings are available within the article and its supplementary information files or from the corresponding author upon reasonable request.

## Acknowledgments

This work was supported by the European Research Council (ERC) under the European Union’s Horizon 2020 research and innovation program (grant agreement no. 677772). We thank Enas Abu-Shah for numerous discussions and critical reading of the manuscript.

## Author contributions

E.N., A.A-S., and J.S. conceived the study. S.B. ran numerical simulations. E.N. and H.S. fabricated the models. L.B-L. guided clinical protocols. E.N. performed experiments. E.N. and A.A-S. analyzed data. E.N. wrote the original draft. E.N., A.A-S., D.W., O.H., M.G., L.B-L., and J.S. discussed results and reviewed the manuscript. E.N., A.A-S., and J.S. revised the final manuscript. J.S. acquired the financial support for the project leading to this publication.

## Competing interests

The authors declare no competing interests.

## Supplementary materials

**Supplementary Figure 6:**
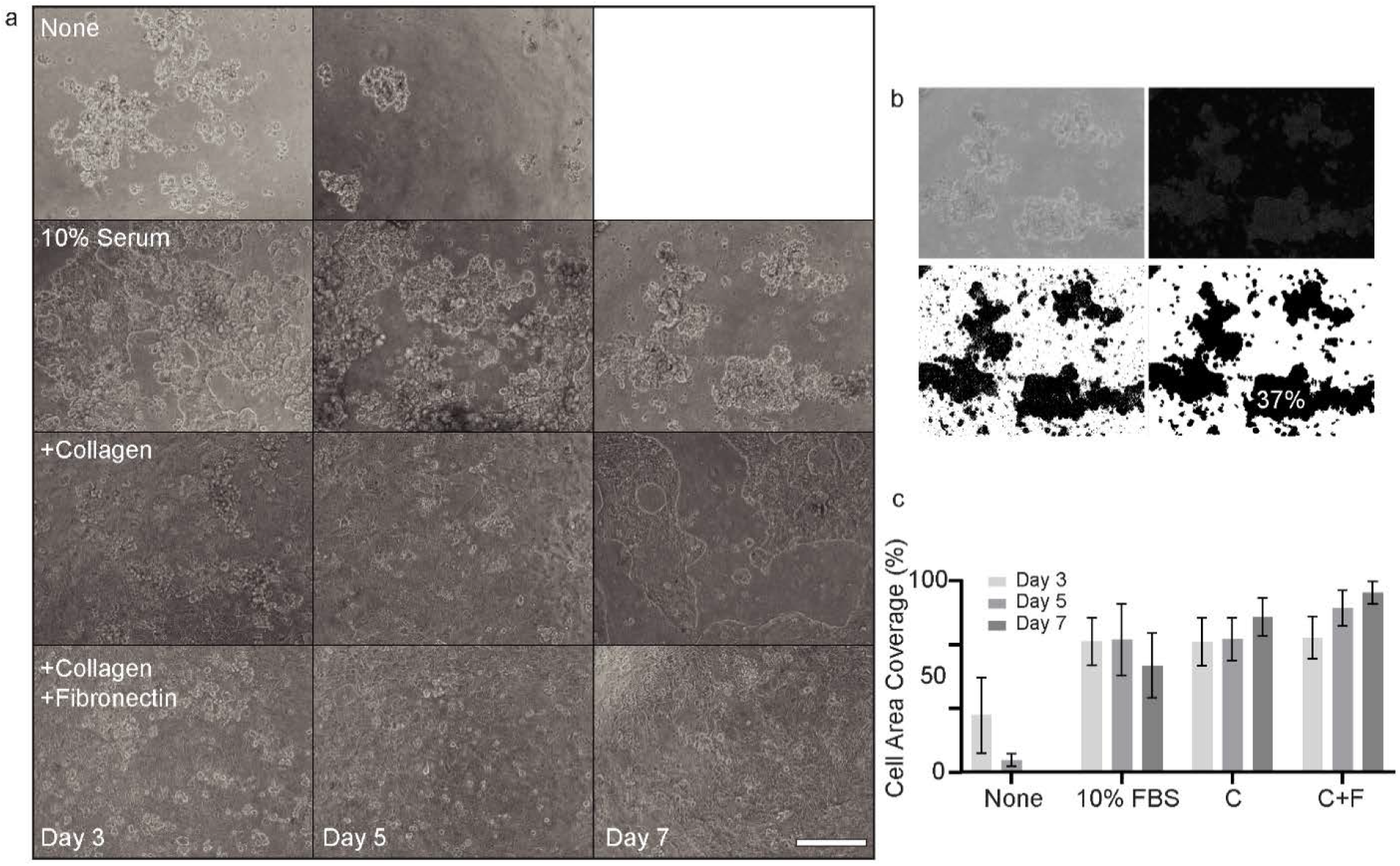
Model lumen coating study for optimal Calu-3 culture conditions. (**a**) Bright-field microscopy images of Calu-3 bronchial epithelial cells inside a six-well plate covered with PDMS at three, five, and seven days following initial seeding directly on a PDMS surface, i.e., no coating (top row), with 10% fetal bovine serum (FBS) (second row), with 1% v/v collagen (third row) and with a combination of 1% v/v collagen and 1% v/v fibronectin (bottom row). No images were taken after seven days with no coating due to the absence of any remaining live cells. (**b**) A representative image is shown undergoing four segmentation processing steps used to quantify the area covered by cells in each image (ImageJ, National Institutes of Health). (**c**) Plot summarizing the results of the study for each coating type at three imaging time points. Error bars signify standard deviation (N=5).

**Supplementary Figure 7:**
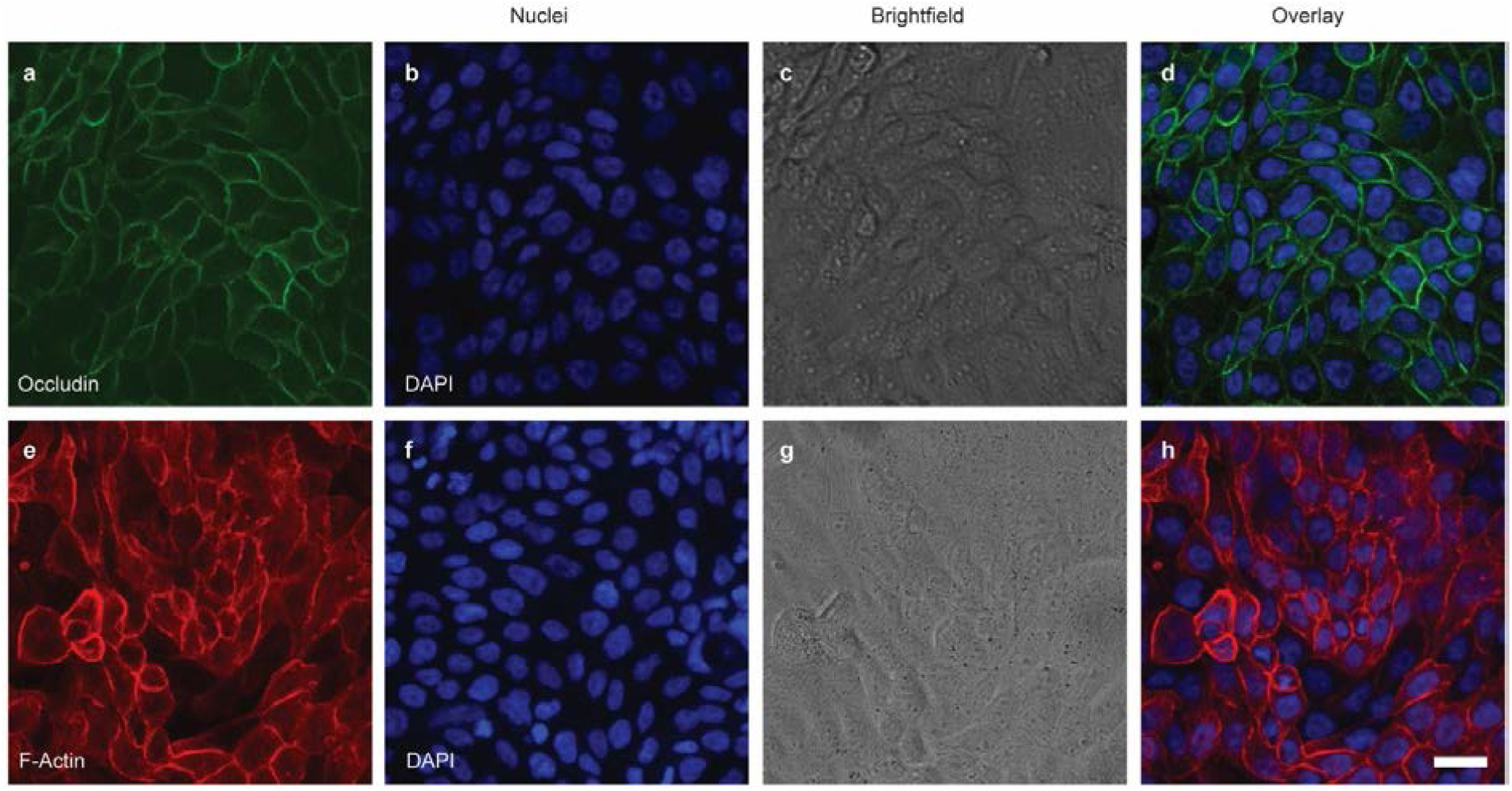
Bronchial epithelial cell line (Calu-3). Stained confocal microscopy images showing the development of a confluent monolayer when cultured on top of collagen and fibronectin-coated surface. Occludin (top left) stained green, an integral protein localized at the epithelial cells’ tight junctions. F-actin (bottom left) stained red, a major component of the cytoskeleton. The second column features nuclei stained with blue DAPI. Third column bright-field microscopy. The last column shows all images as combined overlays characterizing the epithelial monolayer.

**Supplementary Figure 8:**
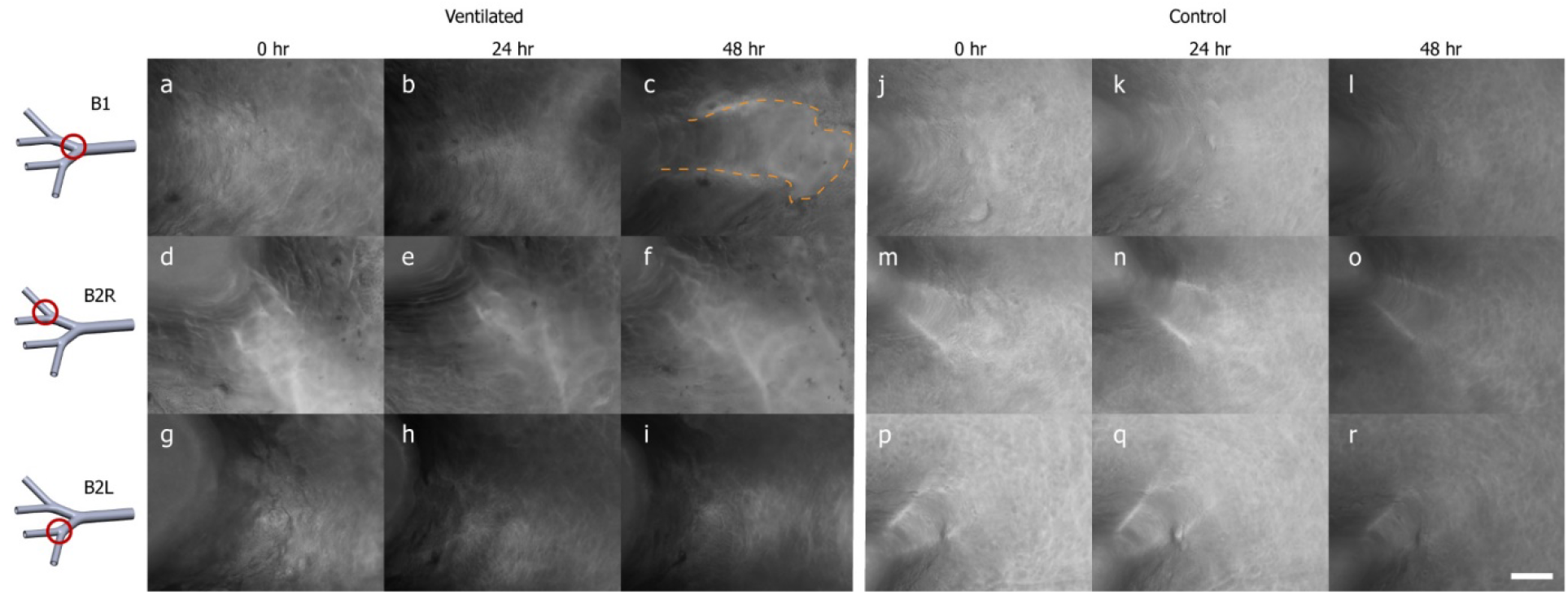
Cell detachment localized in the high-stress region at first bifurcation appears 48 h after ventilation exposure. Bright-field microscopy images of live cells in the same representative ventilated (a-i) and control (j-f) models compare epithelial cell monolayer integrity at three regions of interest and three time points following ventilation exposure. The regions are the three bifurcations of the model labeled B1 for the first bifurcation(top row; a-c, j-l), B2R for the second bifurcation on the right (second row; d-f, m-o), and B2L for the second bifurcation on the left (bottom row; g-i, p-r) are shown at three time points following ventilation exposure: 0 (i.e., immediately after), 24 h and 48 h. Localized cell detachment was seen only in ventilated models and only at the first bifurcation, highlighted with a dashed line in (c) appearing 48 h following exposure. The scale bar for all images is 100 µm.

**Supplementary Figure 9:**
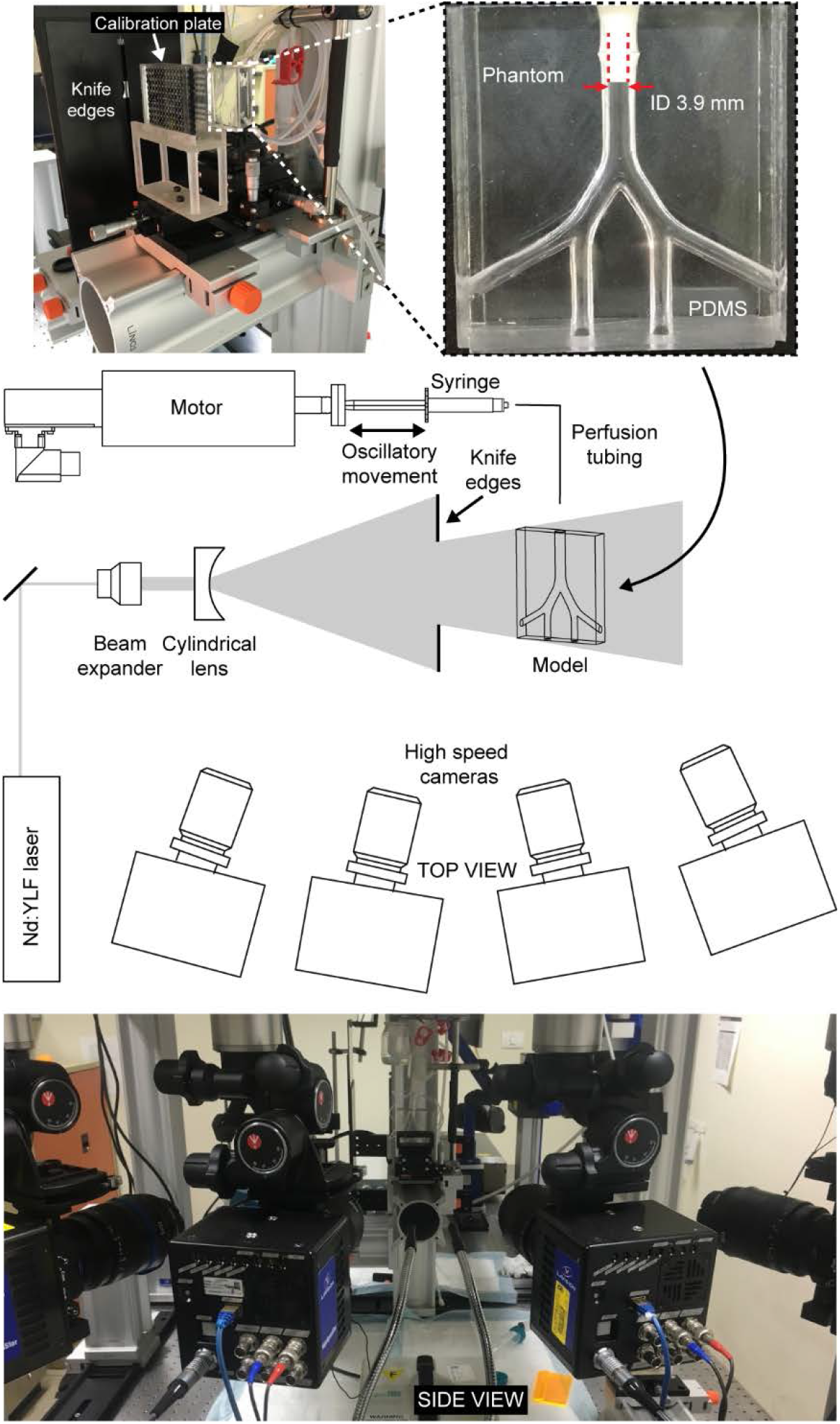
Tomographic particle image velocimetry (TPIV) experimental set-up comprising an Nd:YLF high-powered laser, four high-speed cameras, auxiliary optics, and a phantom PDMS model perfused by a linear motor and syringe. A calibration plate is used for mapping the x-y-z world point to the camera chips, and knife edges provide a well-defined border for uniformly illuminating the measurement volume. Note that in place of the endotracheal tube (ETT) used in the clinically replicated ventilation *in vitro* experiment, a nylon tube to tube connector is used to prevent leaks.

**Supplementary Figure 10:**
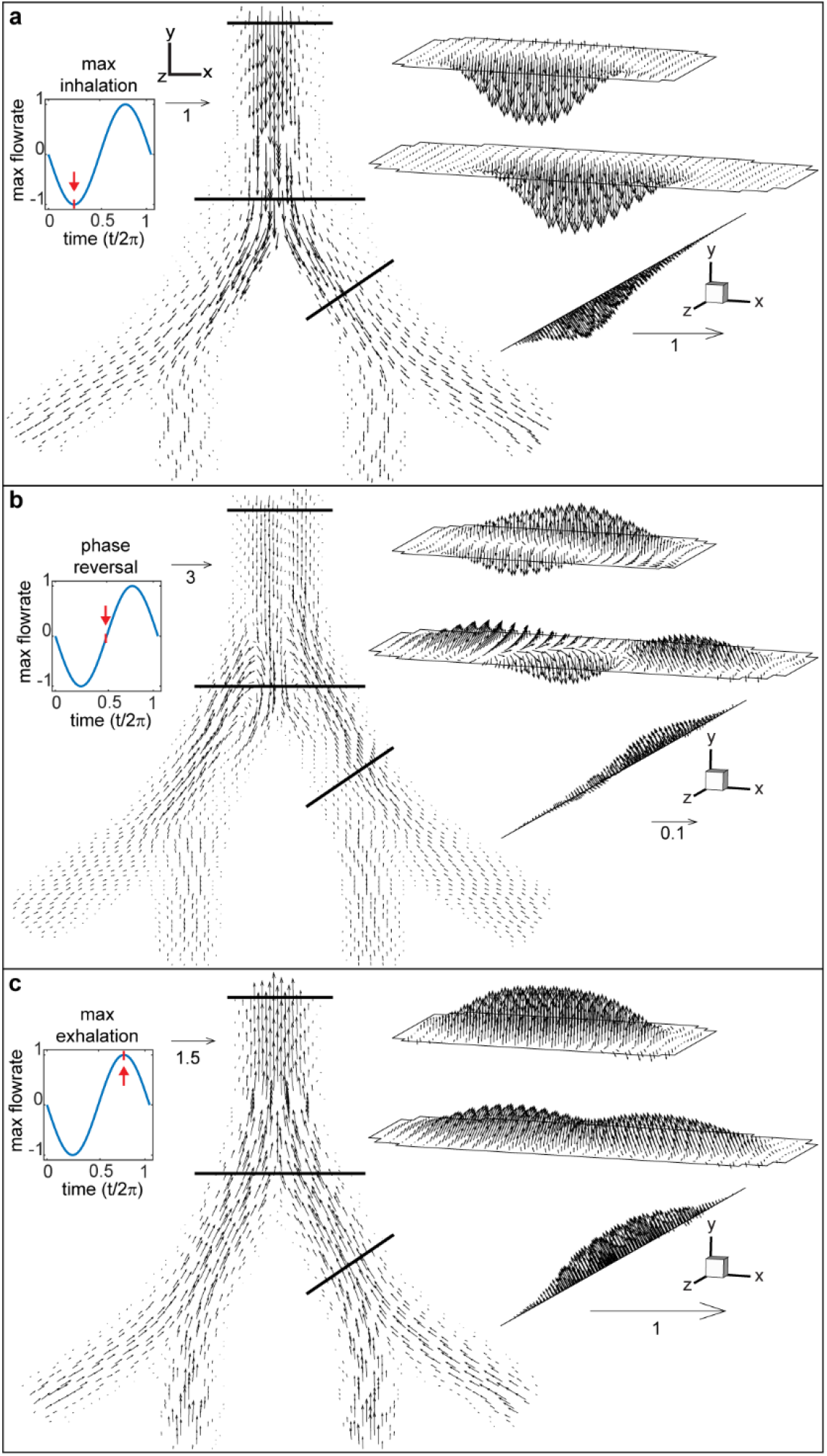
Instantaneous 3D velocity vector fields shown at various time points during a ventilation cycle, measured using tomographic particle image velocimetry (TPIV).

**Supplementary Figure 11:**
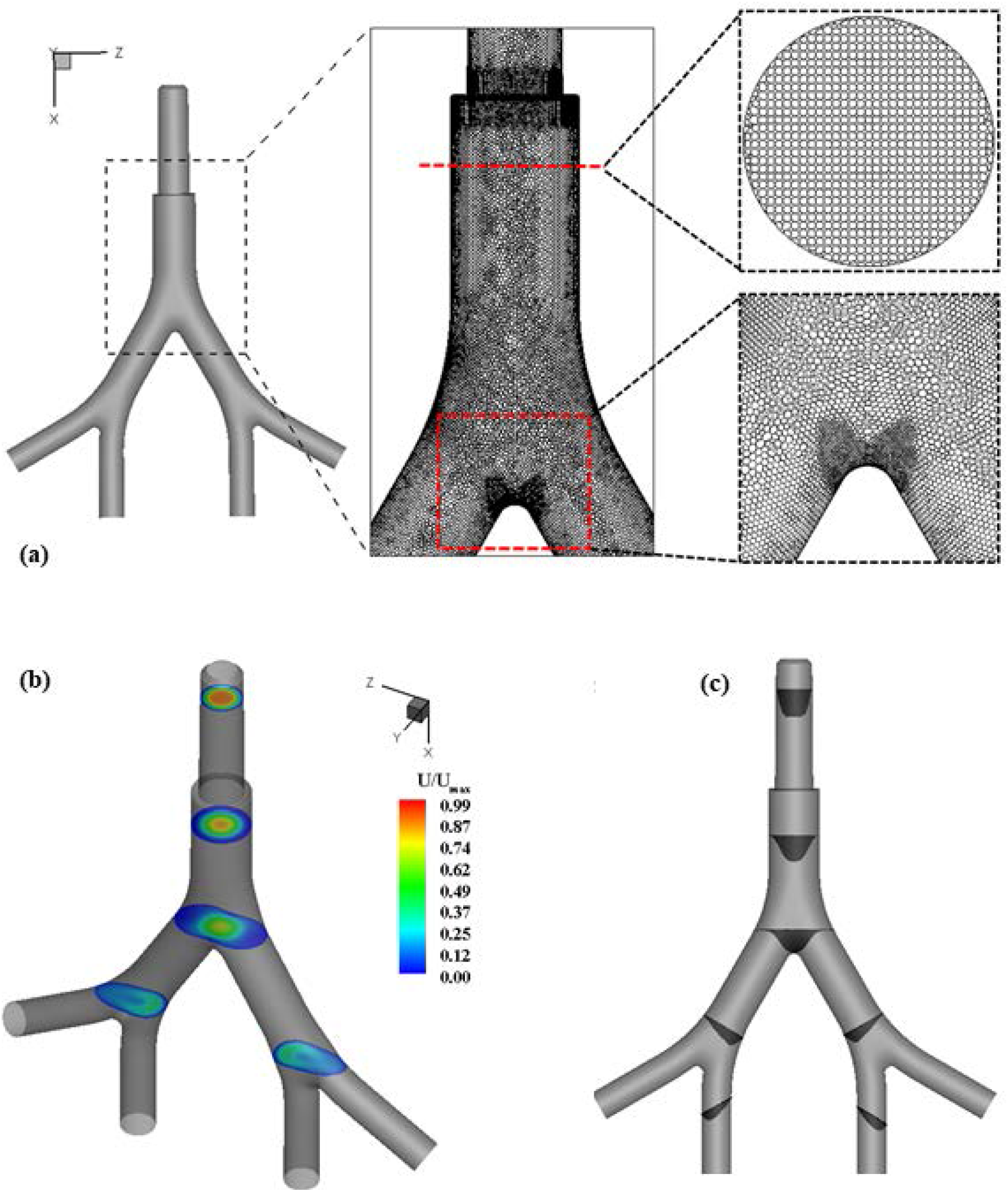
Computational fluid dynamics (CFD) setup. (**a**) The model geometry is meshed with tetrahedral cells in ANSYS ICEM and then transformed into polyhedral meshes in ANSYS Fluent. The mesh is refined at the main carina as well as at the secondary bifurcation zones. In the top enlarged inset, a perpendicular cross-section displays the mesh grid in the upper portion of the trachea, whereas in the bottom inset, an enlarged view of the carina shows increased mesh refinement. In (**b**), the non-dimensional mean velocity contours are plotted at several orthogonally-sliced planes for a representative ventilation case (*α* = 2) and in vector format displayed at the same locations shown in (**c**).

**Supplementary Figure 12:**
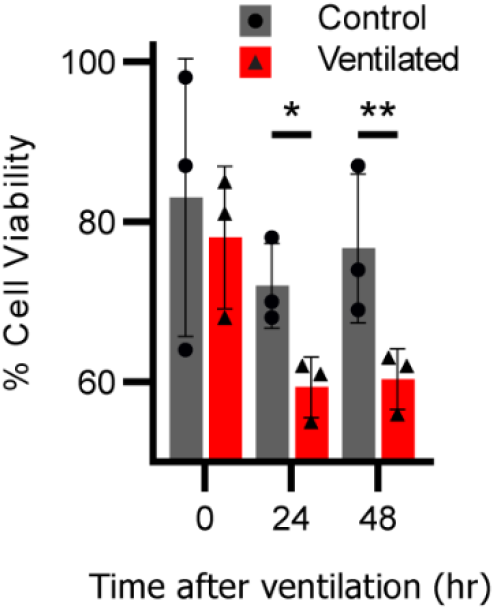
Cell viability assay. Cell viability was measured by 10% alamarBlue-supplemented culture medium for 2 h in models analyzed three times over 48 hours following the ventilation exposure. Immediately after the exposure (i.e., at time 0 hr), near 100% cell viability is measured, indicating metabolic activity has not been affected in either control or ventilated groups. At 24 h and 48 h, cell viability is reduced by ∼40% in the ventilated group while remaining near baseline levels in the control group.

## Notes

### Competing Interest Statement

The authors have declared no competing interest.

### Summary of Updates

This version corrects formatting issues from the previous PDF conversion

